# Drug related and psychopathological symptoms in HIV-positive men who have sex with men who inject drugs during sex (Slamsex): Data from the U-SEX GESIDA 9416 Study

**DOI:** 10.1101/703272

**Authors:** Helen Dolengevich-Segal, Alicia Gonzalez-Baeza, Jorge Valencia, Eulalia Valencia-Ortega, Alfonso Cabello, Maria Jesus Tellez Molina, Maria Jesus Perez Elias, Regino Serrano, Leire Perez Latorre, Luz Martín-Carbonero, Sari Arponen, Jose Sanz Moreno, Sara De La Fuente, Otilia Bisbal, Ignacio Santos, Jose Luis Casado, Jesus Troya, Miguel Cervero Jimenez, Sara Nistal, Guillermo Cuevas, Javier Correas Lauffer, Marta Torrens, Pablo Ryan, on Behalf of the U-SEX GESIDA 9416 Study

## Abstract

**Objectives:** Intravenous sexualized drug use also known as slamsex seems to be increasing among HIV-positive men who have sex with men (MSM). This practice may entail severe consequences for physical and mental health in this population. Research on the subject is scarce. The aim of our study was to describe the psychopathological background of a sample of HIV positive MSM who practiced slamsex during the previous year and compare the physical and psychological symptoms between these participants and those who practiced sexualized drug use (SDU) or chemsex without slamsex.

**Design and Methods:** Participants (HIV-positive MSM) were recruited from the U-Sex study in 22 HIV clinics in Madrid during 2016-17. All participants completed an anonymous cross-sectional survey on sexual behaviour and recreational drug use. The present analysis is based on HIV-positive MSM who had practiced SDU.

**Results:** The survey sample comprised 742 participants. Of all the participants who completed the survey, 216 (29.1%) practiced chemsex, and of these, 34 (15.7%) had practiced slamsex. Participants who practiced slamsex were more likely to have current psychopathology (depression, anxiety and drug related disorders) than chemsex users. In addition, participants who practiced slamsex had more high-risk sexual behaviours, polydrug use and were more often diagnosed with sexually transmitted infections (STIs) and hepatitis C than those who did not inject drugs. Compared with patients who did not inject drugs, patients who engaged in slamsex showed more severe drug related symptoms (withdrawal and dependence), symptoms of severe intoxication (loss of consciousness), and severe psychopathological symptoms related to SDU, such as paranoid thoughts and suicidal behaviour.

**Conclusion:** Slamsex (intravenous SDU) is closely associated with current psychiatric disorders and severe drug-related and psychiatric symptoms.

## BACKGROUND

Chemsex, or sexualized drug use (SDU), was first described in UK as the intentional use of recreational drugs in order to enhance sexual relations between gay, bisexual and other men who have sex with men (MSM), usually for long periods of time and often with multiple partners (1). The main drugs involved in this practice are mephedrone, γ-hydroxybutyrate/γ-butyrolactone (GHB/GBL), and crystal-methamphetamine (Crystal-Meth) (2), although other drugs have been also reported, like ketamine, other synthetic cathinones, 3,4-methylenedioxymethamphetamine (MDMA), cocaine, poppers and erectile-dysfunction drugs, (3). Other aspects of this phenomenon, such as the use of geosocial networking applications to locate or participate in sex parties, should be taken into consideration because of their relevance and implications (4). Intravenous use of psychoactive substances, especially stimulants such as mephedrone and Crystal-Meth in this context, is known as slamming or slamsex (2).

Some studies have suggested that the practice of injecting recreational drugs at sex parties might be increasing among MSM (2,5–7). Both chemsex and slamsex have been described as more prevalent in MSM living with human immunodeficiency virus (HIV-positive) when compared to HIV-negative MSM. A recent UK study of HIV-positive MSM reported that 3% of their sample had injected drugs related to sex in the previous 3 months(8) Similarly, the U-Sex study performed in Madrid showed that 4.5% of the sample had practiced slamsex in the previous year (7).

Slamsex has been associated to sex in group, condomless sex with random partners or fisting practices, which increase the frequency of sexually transmitted infections (STIs) and the transmission of viral infections, such as those caused by HIV and hepatitis C virus (HCV) (9).

Both Mephedrone and Crystal-Meth are potent central nervous system stimulants that also act peripherally. The potency and half-life of mephedrone depends on the route of administration, which varies from an onset of action of half an hour if it is taken orally, with a mild high that can last from 3 to 5 hours, intranasal, with a potent high after 15 minutes and lasting 1-2 hours, and intravenously, with an almost immediate and very potent high with a short duration of 30 to 45 minutes. The rapid onset of action and fast dissipation of effects leads to a compulsive pattern of use and the need to re-dosify almost every hour. Thus, high doses of mephedrone are used in sexual settings, with the consequent risk of overdosing, with altered behaviors and delusive thoughts.

Crystal-Meth is different, as its potency is similar both inhaled or injected intravenously. Either route of administration produces an immediate action of the drug, from 0 to 2 minutes, with a very potent high. If it is injected intravenously, its duration can be quite long, almost 8 hours. Crystal-Meth produces an intense state of excitement, with euphoria, self confidence and sociability. Its withdrawal syndrome is very unpleasant; thus, its addictive potential is very high (10).

Both substances have been related to induced psychotic symptoms in diverse populations (11,12). However, the emergence of psychiatric symptoms in relation to slamsex is scarce, although there is evidence suggesting that mephedrone related to slamsex can induce psychotic symptoms and suicidal conducts (13). Crystal-Meth also has been related to high levels of addiction, psychotic symptoms and other psychiatric disorders in the context of chemsex (14).

Mental health issues have been poorly studied among persons who engage in chemsex and few data are available on the severity of drug-induced symptoms in HIV-positive MSM who practice slamsex.

The aim of our study was to compare the physical and psychological patterns of HIV-positive MSM who practiced slamsex with that of those who practiced chemsex without intravenous injection of drugs. We also explored the presence of psychopathological symptoms and symptoms of substance use disorders induced by drugs in the entire sample, and their correlates. Patients were selected from the U-SEX GESIDA study (7).

### Materials and Methods

The present analysis is nested in the U-SEX GESIDA 9416 study, which was conducted in 22 HIV clinics in the Madrid area from June 2016 to March 2017. This study aimed to calculate the prevalence of chemsex and its associated factors in a sample of HIV-positive MSM in Spain. The inclusion criteria were; age ≥18 years, documented HIV infection and being an MSM. All the participants confirmed to be gay or bisexual. Infectious diseases physicians offered all the participants who met the inclusion criteria the opportunity to participate and gave them a card with a unique code and a link with access to an online survey. The survey was self-completed outside the hospital to ensure anonymity and confidentiality

The online survey was designed “ad hoc” by the research team to evaluate various domains: general sociodemographic data (age, occupational status, income, etc.), HIV infection status (year of diagnosis, treatment, adherence, etc.), sexual behaviours (condom use, receptive anal sex, fisting, etc.), diagnosis of STIs (including HCV), diagnosed psychiatric disorders and history of drug use. If the participant reported any kind of drug use, they were asked if these drugs were used before or during sexual encounters. Chemsex was defined as the intentional use of mephedrone or other cathinones, 3,4-methylenedioxy-N-methylamphetamine (MDMA), methamphetamine, amphetamines GHB/GBL, ketamine, or cocaine during sex. This analysis included participants who reported they had engaged in the practice of chemsex in the last 12 months. The survey evaluated the type of drugs used, the context in which they were used, frequency, route of administration and other aspects referring to the practice of chemsex.

In order to collect psychiatric disorders data, the survey asked general questions regarding previously diagnosed psychiatric disorders and specific questions about “past” or “current” psychiatric disorders diagnosed by a mental health specialist. To conduct the present analysis, we only considered self-reported current psychiatric disorders (diagnosed in the previous year), namely, depression, anxiety, personality, psychosis, and drug-related disorders. Because the survey was self-completed, we used the term “self-reported current psychiatric disorder”.

All participants were asked about dependence, withdrawal, and psychopathological symptoms related to the drugs used in chemsex sessions. To determine drug dependence symptoms, the survey asked about the following items: drugs used more often or in a higher quantity than planned, severe craving, not fulfilling obligations because of drug use, continuing drug use (even when this lead to physical or psychological discomfort), need to increase doses to obtain the same effect and less positive effects with same doses. The presence of 3 or more symptoms of drug dependence during the previous year were considered in the analysis.

In order to collect data on symptoms of withdrawal we asked about the following: severe craving, need to take medications/other drugs to compensate for discomfort, sleep disturbances (insomnia, hypersomnia), agitation, depressive thoughts/feelings, paranoid ideation, suicidal thoughts, suicide attempts, and the need to see a doctor for treatment of discomfort. The presence of 3 or more symptoms of withdrawal/abstinence during the last year were included in the analysis.

Finally, intoxication-related symptoms were assessed based on the following: sleep disturbances, “things done to me that I would not have consented to without being on drugs”, “more sexual risk practices that I don’t do when not on drugs”, unpleasant physical feelings under the effects of drugs, anxiety/panic attacks, irritability, and aggressiveness. Psychotic symptoms (mainly paranoid ideation), loss of consciousness, suicidal thoughts and suicide attempts were considered severe intoxication symptoms.

Details of the study procedures have been previously published (7). In the present study, to clarify the terminology applied when comparing participants, we used the following terms: participants who engaged in slamsex when the SDU was intravenous and participants who engaged in chemsex when the drugs were not consumed intravenously.

The study protocol was approved by the Ethics Committee of Hospital Universitario Gregorio Marañón (HUIL 1606 96/16) and fulfilled the principles of the Declaration of Helsinki (2008).

Study data were collected and managed using the data capture tool Research Electronic Data Capture (REDCap) (15) hosted at “Asociación Ideas for Health”.

### Statistical Analysis

Categorical variables were expressed as absolute and relative frequencies; continuous variables were expressed as median (IQR). Baseline characteristics were compared between participants who had engaged in slamsex and participants who had engage in chemsex during the previous year, using the chi-square test for categorical variables and the *t* test for continuous variables. Variables included in the comparisons were sociodemographic variables, self-reported current psychiatric disorders, physical and severe psychopathological symptoms related to drug use/abuse, sexual behaviors, and medical variables such as, time since HIV diagnosis, self-reported adherence to antiretroviral therapy or STDs diagnosis.

We conducted a logistic regression analysis to explore the association between slamsex and both symptoms of drug use disorders and severe psychopathological symptoms. We separately tested the association of slamsex with the presence of withdrawal (three or more withdrawal symptoms), dependence (three or more dependence-related symptoms), craving (strong need for consumption), paranoid ideation (during or after drug use), suicidal behaviors (suicidal ideation and suicide attempts during or after drug use) and loss of consciousness (during or after drug use).

The univariate analysis was conducted separately to evaluate the association between symptoms of drug-related disorders or severe psychopathological symptoms in the context of chemsex and, other drug-related variables or self-referred psychiatric current disorders. The dependent variables included withdrawal symptoms, severe craving, psychotic paranoid ideation, suicidal behaviours, and loss of consciousness. Independent variables were categorized as the presence/absence of self-referred active depression, self-referred active anxiety, polydrug use (three or more drugs used each time), cathinone use during the previous year, ketamine use during the previous year, GHB use during the previous year and inhaled Crystal-Meth use during the previous year. Thereafter, bivariate logistic regressions were conducted to explore associations regardless of the presence of slamsex. The presence/absence of slamsex was included in the bivariate regression as an independent variable. Independent variables were included in the bivariate analysis only if their p value was ≤.10 in the univariate analysis.

## RESULTS

### 1.1. Baseline characteristics and comparison between slamsex and chemsex

Of a total of 742 HIV-positive MSM who completed valid surveys in the U-Sex Study, the present analysis included all the participants who had engaged in chemsex during the previous year (N=216). Participants in our sample were mainly Spanish born (71.3%), middle aged (median=38; IQR: 33-44), and with a university education (63.9%). In addition, 70.8% had a salary of more than 1000 euros per month, and 42% were in a stable relationship. The median years with HIV diagnosis was 5 years (IQR: 2-11). More than 90% were receiving antiretroviral therapy and of these, 3% reported having taken less than 90% of doses (poor adherence). In our sample, thirty-four participants (15.7%) had practiced slamsex during the previous year. A comparison with HIV-positive MSM who did not engage in chemsex in our sample has been reported elsewhere (7).

When participants who had engaged in slamsex during the previous year were compared with those who engaged in chemsex, no differences were found regarding sociodemographic or medical variables. Compared with people who engaged in chemsex, people who had engaged in slamsex were less likely to have a stable partner (26.5 vs. 45.6%, *P*=.039) and tended to have more frequently poor adherence to antiretroviral therapy (9.1 vs. 1.9%, *P*=.061).

Comparisons based on the type of drug used in both groups of participants are shown in table 1. Participants who engaged in slamsex had higher rates of polydrug use (3 or more drugs per session), use of mephedrone and other cathinones, Crystal-Meth, ketamine, and intrarectal use of drugs. They also had higher rates of high-risk drug use behaviours, such as sharing needles or other drug paraphernalia. Symptoms related to drug abuse/dependence and severe psychopathological symptoms associated with the practice of slamsex and chemsex are shown in table 2. Regarding the kind of drugs injected intravenously, the most frequent were mephedrone or other cathinones (94.1%), then ketamine (17.6%), Crystal-Meth (5.9%) and cocaine (5.9%).

**Table 1.** Comparisons between Participants who practiced Chemsex and who practiced Chemsex in terms of type of drug used in the previous year.

|  | **Entire sample**<br>**(N=216)** | **Chemsex**<br>**(n=182)** | **Slamsex**<br>**(n=34)** | **P value** |
| --- | --- | --- | --- | --- |
| **Polydrug No. (%)** | 98(45.4) | 70(38.5) | 28(82.4) | .000 |
| **Poppers No. (%)** | 170 (78.7) | 140 (76.9) | 30 (88.2) | .139 |
| **Mephedrone or other cathinones**<br>**No. (%)** | 150 (69.4) | 116 (63.7) | 34 (100) | .000 |
| **Cocaine No. (%)** | 171 (79.1) | 146 (80.2) | 25 (73.5) | .378 |
| **MDMA No. (%)** | 105 (48.6) | 87 (47.8) | 18 (52.9) | .582 |
| **GHB No. (%)** | 155 (71.7) | 128 (70.3) | 27 (79.4) | .280 |
| **Crystal methamphetamine No.**<br>**(%)** | 64 (29.6) | 47 (25.8) | 17 (50) | .005 |
| **Ketamine No. (%)** | 78 (36.1) | 57 (31.3) | 21 (61.8) | .001 |
| **Rectal use of drugs No. (%)** | 44(20.4) | 24(13.2) | 20(58.8) | .000 |
| **High-risk drug use No. (%)** | 168 (77.8) | 135 (74.2) | 33 (97.1) | .003 |

**Table 2.** Self-reported psychiatric symptoms during or after chemsex and slamsex.

|  | **Entire sample**<br>**(N=216)** | **Chemsex**<br>**(n=182)** | **Slamsex (n=34)** | **P**<br>**value** |
| --- | --- | --- | --- | --- |
| **3 or more dependence symptoms**<br>**No. (%)** | 60(27.8) | 40(22) | 20(58.8) | .000 |
| **3 or more withdrawal symptoms**<br>**No. (%)** | 98(45.8) | 72 (39.6) | 26 (76.5) | .000 |
| **Intense craving. No. (%)** | 55 (25.5) | 34 (18.5) | 21 (61.8) | .000 |
| **Interference with work, social or**<br>**family life No. (%)** | 68 (31.5) | 46 (25.3) | 22 (64.7) | .000 |
| **Paranoid ideation No. (%)** | **30 (15.3)** | **20 (11)** | **10 (29.4)** | **.004** |
| **Suicidal ideation No. (%)** | 33 (15.3) | 22 (12.1) | 11 (32.4) | .003 |
| **Suicide attempt No. (%)** | 30 (13.8) | 19 (10.4) | 11 (32.4) | .001 |
| **Loss of consciousness. No. (%)** | 33 (15.3) | 23 (12.6) | 10 (29.4) | .013 |

Participants who engaged in slamsex showed a significantly higher percentage of sexual risk behaviours than those who practiced chemsex, as follows: fisting (73.5 vs. 38.5%, *P*=.001), fisting without a glove (67.7 vs. 28%, *P*=.001), condom use in less than half of sexual relations (93.1 vs. 48.3%, *P*=.001) and more than 20 sexual partners in the previous 6 months (70 vs. 39.6%, *P*=.002). As for STDs, people who had engage in slamsex more often had gonorrhea (43.4 vs. 61.8%, *P*=.049), syphilis (62.6 vs. 88.2%, *P*=.004) and hepatitis C (18.1 vs. 61.8%, *P*=.000) than people who engage in chemsex.

A self-reported current psychiatric disorder was more common among participants who engaged in slamsex than in those who engaged in chemsex, with the conditions reported as follows: depressive disorder (61.8 vs. 28%, *P*=.0001), anxiety disorder (47.1 vs. 23.1%, *P*=.004), and drug use disorders (drug-dependence) (38.2 vs. 15.4%, *P*=.002).

### 1.2. Correlates of severe physical and psychopathological symptoms related to drug use

The simple logistic regression conducted to explore the association between slamsex and the presence of symptoms of drug use disorders or severe psychopathological symptoms related to drug use revealed a significant association. Compared with participants who had engaged in chemsex,, those who engaged in slamsex were five times more likely to had experienced withdrawal symptoms (OR: 4.97 [2.13-11.57], *P*=.0001) and seven times more likely to had experienced intense craving (OR: 7.03 [3.21-15.43], *P*=.0001). Moreover, during or after drug use they were three times more likely to experience suicidal ideation (OR: 3.48 [1.48-8.10], *P*=.004), psychotic paranoid ideation (OR: 3.38 [1.41-8.07], *P*=.006) and loss of consciousness (OR: 2.88 [1.22-6.79], *P*=.016)..

Figure 1 shows the associations between other drug-related variables or current self-reported psychiatric diagnosis and, the presence of symptoms of drug related disorders and severe physical and psychopathological symptoms related to drug use (suicidal ideation, paranoid ideation and loss of consciousness), regardless of the presence of slamsex. Patients who self-reported current depressive disorders more frequently had withdrawal symptoms. Active anxiety, cathinone use and GHB use were also associated with the presence of withdrawal symptoms. Moreover, participants who inhaled Crystal Meth more frequently experienced severe craving, and, those who inhaled Crystal-Meth or used multiple drugs were significantly more likely to present symptoms of drug-dependence. Suicidal ideation was only associated with self-reported depression and anxiety disorders; paranoid ideation was associated with anxiety disorders, polydrug use, and inhaled Crystal-Meth. Finally, loss of consciousness was related to polydrug use, GHB use, ketamine use, and inhaled Crystal-Meth (Fig 1).

## DISCUSSION

The present study provides novel findings regarding the slamsex phenomenon in a sample of HIV-infected MSM who engage in SDU. In our sample, 216 subjects engaged in chemsex. From this sub-sample, 34 subjects (15.7%) engaged in slamsex during the previous year. Compared with those who did not inject drugs, people who had engaged in slamsex more frequently reported high risk sexual behaviours, had more frequently been diagnosed with an STD, and had more frequently reported a current diagnosis of a psychiatric disorder. In addition, compared with participants who engaged in chemsex, participants who had engaged in slamsex in the previous year had more drug-related adverse effects such as symptoms of withdrawal and dependence or severe physical and psychopathological symptoms such as psychotic paranoid ideation, suicidal behaviours and loss of consciousness.

There is some research regarding the prevalence of slamsex and associated high risk behaviours among MSM. The Unlinked and Anonymous Monitoring (UAM) survey of people who inject drugs reported that since 2000, the proportion of MSM who inject drugs has increased significantly (4.4% in 2000/2001 to 8.1% in 2014/2015, *P*<0.001). They also reported the presence of higher-risk behaviours associated with injecting such as needle/syringe sharing (15% vs 11%, *P*=0.07) and having more than 10 sexual partners among MSM who injected drugs than among MSM who did not inject drugs (25% vs. 4.0%, *P*<0.001) (6). A recent study from an Australian cohort of MSM reported a high life prevalence of injecting drugs (10.3%); the prevalence of injection in the previous six months was 4.7% in this population. The authors reported that injecting drugs was associated with high-risk sexual practices such as having multiple sex partners, group sex with casual partners and condomless anal intercourse with casual partners (16). In the case of HIV-positive MSM, the ASTRA study (17) reported that of 2248 HIV-positive sexually active MSM recruited in 2011-2012, 1138 (51%) had used recreational drugs in the previous three months and the prevalence of injection drug use was 3% (n=68). The Positive Voices Study reported that 105 of 392 sexually active HIV-positive MSM (29%) had engaged in chemsex during the previous year. Among these, the prevalence of slamsex was 33.3% (18). The prevalence of slamsex in our study could be directly compared with the findings of the Positive Voice study only because of methodological and sample similarities. While the rate of chemsex reported is similar to the rate we report previously (29%) in the U-Sex Study (7), the authors found higher rates of slamsex among their participants (33.3 vs. 15.6%). Therefore, we think that regional differences in slamsex frequencies should be explored in future studies. Otherwise, the most dangerous profiles of drug use and sexual practices found in the above mentioned studies in samples of MSM who injected drugs are congruent with the higher rates of polydrug use, rectal use of drugs, sharing drug paraphernalia, sexual risk behaviours and STDs found among those who engaged in slamsex in the present study.

We found that the most common intravenous drugs used during slamsex were mephedrone or other synthetic cathinones (94.1%), followed by ketamine (17.6%), Crystal-Meth (5.9%) and cocaine (5.9%). To our knowledge, only a few reports discuss the type of drug used by HIV-positive MSM during slamsex. The UAM Survey found high frequencies of injected mephedrone and ketamine among MSM who injected drugs (12% and 9.3%, respectively) (6). Furthermore, data from Antidote, a specialist drug clinic aimed at the gay community in London, UK, showed that 75% of patients used mephedrone in the chemsex context and of these, 80% injected the drug. Of this 80%, 75% were HIV-positive and 70% reported sharing needles (19). The recently published FLUX study, a survey performed in Australian gay and bisexual men, found that of the 1995 respondents, 206 (10.3%) reported having injected drugs and 93 (4.7%) had injected recently, most commonly Crystal-Meth (91.4%) and speed (9.7%), as well as cocaine and ketamine, albeit in low percentages (16). Together with the data reported above, our results suggest that the type of drugs injected in the chemsex context are similar but that there may be regional differences. Drug use in the context of chemsex and slamsex may be changing continuously, as a result of travel by MSM to different countries for leisure, socialization and clubbing and to expand sexual experiences.

The participants in our sample who engaged in slamsex presented higher rates of drug use related adverse symptoms than those who engaged in chemsex. Severe craving and other withdrawal symptoms were more frequent, as was loss of consciousness. The participants also showed higher rates of severe psychopathological symptoms such as paranoid ideation and suicidal ideation or attempts.

Mephedrone and other synthetic cathinones were the main drugs “slammed” in our sample, both as stimulants and as sexual enhancers. The intravenous use of mephedrone has been related to compulsive use, intense craving, binging behaviours and withdrawal symptoms (20). Diverse psychotic symptoms, mainly paranoid ideation, have also been reported for mephedrone, especially if it is consumed intravenously (21,22). In the context of slamming, one case in Spain has been reported in a young HIV-positive man, who experienced persistent mephedrone-induced paranoid delusions, intense anxiety and visual and kinaesthetic hallucinations (13).

Ketamine, cocaine and Crystal-Meth were also consumed in slamsex in our sample, albeit at a lower frequency than cathinones. Injected Crystal-Meth has the potential to induce psychotic symptoms and has been related to drug-related disorders such as abuse or dependence. In slamsex, its potent stimulant effect has been related to high-risk sexual behaviors, with an increased risk of infection by HIV or other STDs (10).

Traditionally, more frequent drug dependence and psychiatric symptoms have been described during intoxication by or abstinence from some drugs if they are used intravenously. Our novel data together with the few previously published findings support the addictive potential and severe psychopathological consequences of drugs injected in the chemsex context.

Other variables related to drug use might modulate the severity of physical and psychopathological symptoms induced by drugs in the context of SDU. Regardless of the presence of slamsex, use of inhaled Crystal-Meth, GHB use (oral), ketamine use, polydrug use, and self-reported depression and anxiety disorders were associated with more severe physical and psychopatological symptoms related to drug use in our sample of HIV-infected MSM who engaged in chemsex. In particular, inhaled Crystal-Meth was associated with higher rates of drug dependence and withdrawal symptoms. Moreover, participants who used inhaled Crystal-Meth more frequently had psychotic paranoid ideation and loss of consciousness experienced during or after drug use.

In addition to intravenous injection, inhaled Crystal-Meth has been used by MSM at sex parties for quite some time. The potent disinhibiting effect of this drug has been related to high-risk sexual behaviours and an increase in the frequency of STIs, particularly HIV infection(23). Furthermore, drug-dependence has been described in MSM who inject Crystal-Meth and who are also more prone to comorbid psychiatric disorders and suicidal behaviour (10). Induced psychotic symptoms have been reported in other populations (24), although other psychopathological symptoms induced by inhaled Crystal-Meth in chemsex are scarcely known.

Loss of consciousness was also associated with GHB and ketamine use in our sample. In addition, participants who used more than 3 drugs (polydrug use) had higher rates of loss of consciousness and paranoid ideation and tended to show more pronounced symptoms of drug dependence. This observation must be taken into account, because GHB is usually consumed in combination with other drugs. GHB is frequently related to loss of consciousness, owing to its depressive effect on the central nervous system and because it accumulates over time (2). In addition, the combination of GHB with mephedrone, Crystal-Meth and alcohol increases the risk of drug-drug interactions and overdose, with loss of consciousness and respiratory depression (2). Although ketamine is a dissociative anaesthetic that acts as a stimulant at low doses, with higher doses, polydrug use and intravenous injection, it can increase the risk of loss of consciousness and cardiovascular toxicity in recreational settings (25) such as chemsex, as reported in the present study. Our results are congruent with the effects of these drugs (Crystal-Meth, GHB, mephedrone) previously known. In our opinion this results help to understand the role of each type of drug and route of administration in the severe consequences that may be experienced by some people engaged in chemsex.

Finally, self-reported current diagnosed psychiatric disorders may have played a significant role among the chemsex users in our sample. Regardless of the presence of slamsex, those participants who self-reported current depression more frequently experienced withdrawal symptoms and suicidal ideation during or after drug use. Participants with current anxiety disorders also reported higher rates of withdrawal symptoms, suicidal ideation and paranoid ideation in this context. Moreover, participants who engaged in slamsex were more likely to have anxiety and depression.

While there is evidence that HIV-positive MSM frequently present mental health problems such as depression, anxiety, suicidal behaviour and drug-related disorders, there is little research on the effect of these variables on the health consequences of chemsex practice in this population. The initial published data suggest that HIV-positive MSM who practice chemsex had a higher frequency of depression and anxiety disorders than HIV-positive MSM who did not (8). Other studies on chemsex did not report psychopathological diagnoses, but rather analysed emotional distress and psychological discomfort associated with chemsex. It has been suggested that some vulnerability factors related to problematic chemsex may be the so-called “minority stressors” such as negative internalised homophobia, fear of disapproval, experience of discrimination and a negative self-concept (26).

Therefore, according to the syndemic approach, mental health disorders in HIV-infected MSM appear to increase vulnerability to develop drug abuse disorders and sexual risk behaviours, acting in a syndemic framework by which disease outcomes and the social conditions that contribute to their proliferation sustain each other (26). Consequently, a multidisciplinary approach is necessary to address the situation appropriately. Although our data do not enable us to speculate on causality, in our opinion, the presence of depression and anxiety among HIV-positive MSM who engage in slamsex could indicate vulnerability to develop more severe physical and psychopathological consequences. Moreover, people with previous mental health problems may be more likely to start chemsex and become involved in high-risk practices such as slamsex. We also found that suicidal behaviour in slamsex users was associated with reported current depression and anxiety. These findings can be interpreted in two ways: first, intravenous use of particular drugs such as synthetic cathinones or other stimulants can trigger suicidal ideation in vulnerable subjects; second, subjects with current depression or anxiety may be more prone to use drugs intravenously. The presence of psychopathology along with intravenous drug use can lead to suicidal ideation and suicide attempts, as well as psychotic symptoms. More research is needed to know the causality and interaction between these variables in people who had engaged in chemsex and slamsex.

We think it is important to evaluate the mental health of HIV-positive MSM alongside other routine evaluations conducted in HIV clinics. The detection of psychiatric disorders and their appropriate treatment can prevent other mental and physical consequences of drug use in this population. In addition, approaches such as reducing the harm caused by drug-use can be more effective in people who are not willing to stop drug use in relation to sex. It is necessary to create multidisciplinary approaches in the prevention and treatment of the consequences of chemsex.

To our knowledge we provide for the first time a detailed analysis about drug-related and severe psychopathological symptoms experienced in people engaged in slamsex. We also report data that that increases knowledge about the role of different types of drugs, routes of consumption and psychiatric disorders in drug addiction and psychopatological consequences of chemsex practices among HIV-positive MSM.

Our study is subject to the limitations inherent to cross-sectional survey-based studies, especially response bias. Although we used limited time periods in questions that depended on memory, recall bias could distort the accuracy of the results. Furthermore, we were unable to confirm causality because of the cross-sectional nature of the study. Further longitudinal studies should be performed to compare our results in order to be able to confirm that slamsex can be related to previous psychopathology and may have drug-related and severe psychopathological symptoms in HIV-positive MSM. Another limitation is that the psychiatric diagnosis or drug-related symptoms were self-reported. Although the questionnaire specified previous or current diagnosed psychiatric disorders diagnosed by a psychiatrist or other mental health specialist, the survey did not have standardized diagnostic scales. The exploratory nature of this study led to the inclusion of a large number of variables and the “ad hoc” design of the survey, using sometimes particular slang of the phenomenon in Spain. However, questions about substance dependence and whithrawal were elaborated following DSM-IV-rev criteria. Future studies should include standardized screening scales for mental disorders, substance use disorders, presence of craving or specific psychopatological symptomatology to allow a detailed measurement of specific variables.

Our results suggest that slamsex is relatively common, although it does not appear to be generalized among HIV-positive MSM who practice chemsex in Spain. People who engage in slamsex appear to have high-risk practices associated with both drug use and sexual behaviour in comparison with people who engage in chemsex. Also, people who engage in slamsex, are more likely to experience drug-related induced psychopathological symptoms and symptoms of drug dependence. Moreover, the non-injected use of other substance such as Crystal-Meth, GHB/GBL or ketamine and the presence of psychiatric disorders might also contribute to severe consequences for the physical and mental health of persons who engage in chemsex.

## ACKNOWLEDGMENTS

The authors thank the study patients for their participation, Thomas O’Boyle for writing assistance during the preparation and Ideas for Health Association and Fundación SEIMC-GESIDA for the study management.

Thomas O’Boyle for writing assistance during the preparation

## Contributors

The members of the U-SEX GESIDA 9416 Study Group are listed by centre below; P. Ryan, J. Troya, G. Cuevas, Hospital Universitario Infanta Leonor, Madrid, Spain; J. González-García, J.I. Bernardino, V. Hontañón, I. Pérez-Valero, A. Gonzalez, Hospital Universitario La Paz, Madrid, Spain; S. Moreno, M.J. Pérez, J.L. Casado, A. Moreno, M. Sanchez-Conde, M.J. Vivancos, C. Gómez, S. Serrano, Hospital Universitario Ramón y Cajal, Madrid, Spain; J. Berenguer, A. Carrero, L. Perez-Latorre, T. Aldamiz, Hospital Universitario Gregorio Marañón, Madrid, Spain; A. Cabello, Hospital Fundacion Jimenez Diaz, Madrid, Spain; M. Cervero. Hospital Universitario Severo-Ochoa, Madrid, Spain; S. Arponen, A. Gimeno, C. Montero Hospital de Torrejón, Madrid, Spain; S. Nistal, Hospital Rey Juan Carlos, Madrid, Spain; J. Valencia, J.Gutiérrez, A. Morro, Madrid Positivo, Madrid, Spain; I. Suarez, E. Malmierca, Hospital Universitario Infanta Sofía, Madrid, Spain; J. Sanz, Hospital Universitario de Alcalá, Madrid, Spain; I. Terrancle, R. Monsalvo, Hospital Universitario del Tajo, Madrid, Spain; O. Bisbal, M. Matarranz, M. La Torre, R. Rubio, L. Dominguez, Hospital Universitario 12 de Octubre, Madrid, Spain; R. Serrano, P. Sanz, H. Dolengevich, Hospital Universitario de Henares, Madrid, Spain; A. Diaz, S. de la Fuente, A. Moreno, Hospital Universitario Puerta de Hierro, Madrid, Spain; J.A. Melero, Hospital Universitario Infanta Cristina, Madrid, Spain; M.J. Tellez, V. Estrada, J. Hospital Universitario Clinico San Carlos, Madrid, Spain; M. Estebanez, Hospital Universitario Gomez-Ulla, Madrid, Spain; A. Gomez, I. Santos, L. García-Fraile, J. Sanz-Sanz, C. Sarriá, A. Salas, C. Sáez, Á. Gutiérrez-Liarte, Hospital Universitario La Princesa, Madrid, Spain; T. García-Benayas, T. Fernandez, R. Peñalver, Hospital Universitario del Sureste (Arganda), Madrid, Spain; J.E. Losa, Hospital Universitario Fundacion Alcorcon, Madrid, Spain; H. Esteban, M. Yllescas, Fundación SEIMC-GESIDA, Madrid, Spain.

## Funding/Support

This study was supported by GESIDA.

---

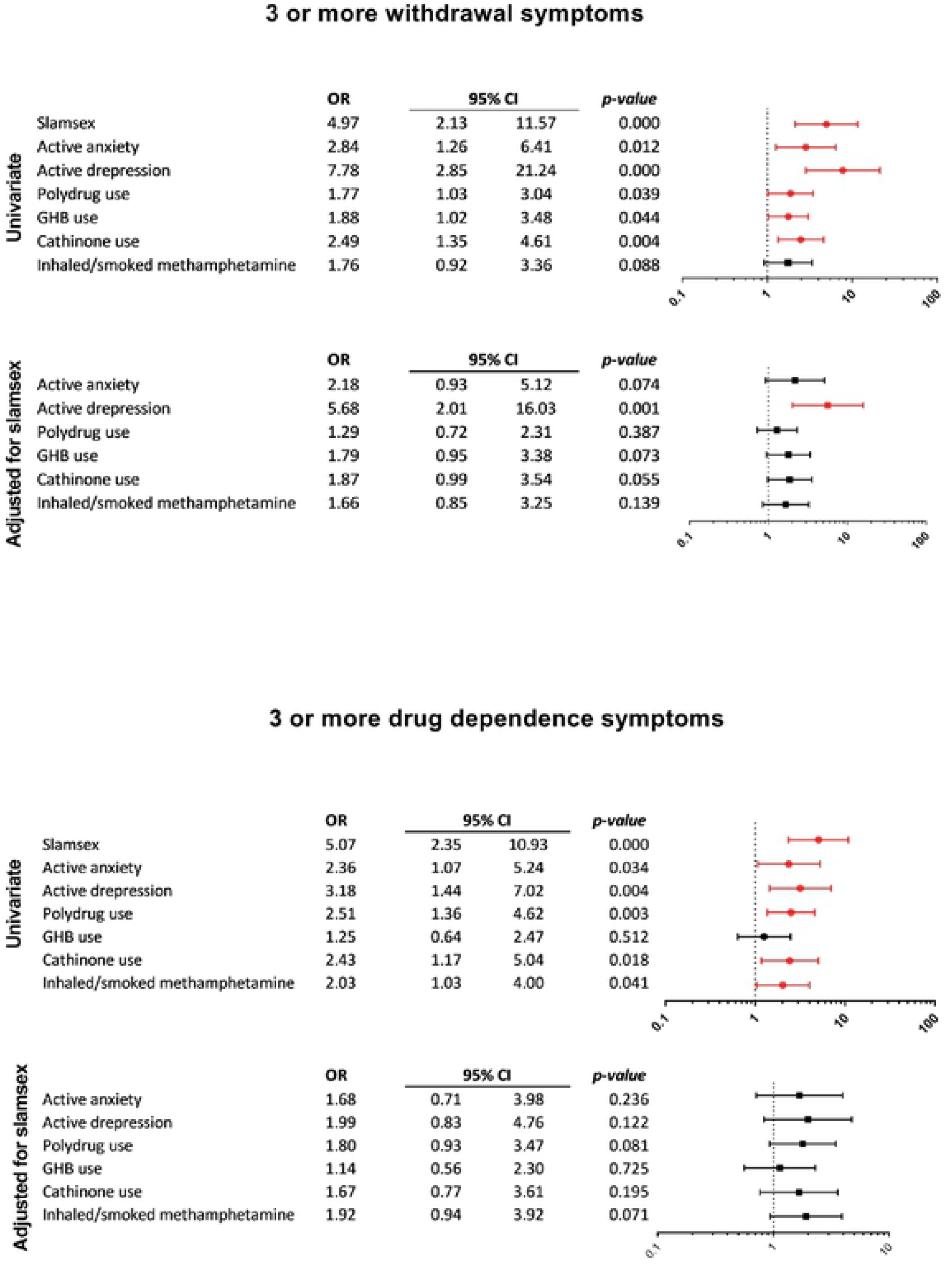

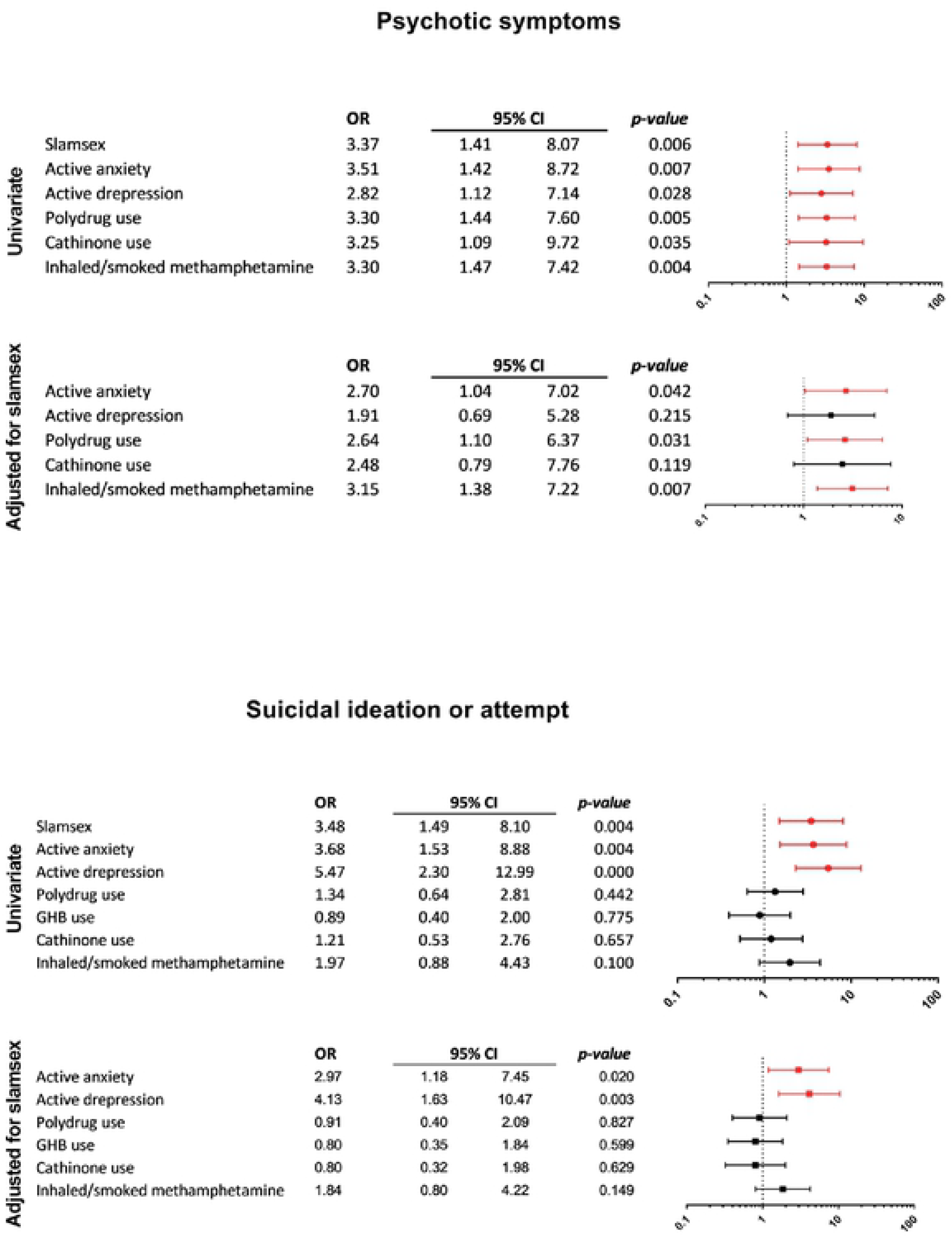

